# A microfluidic platform for *in situ* studies of bacteria electroporation

**DOI:** 10.1101/2024.05.24.595686

**Authors:** Ivan L. Volkov, Zahra Khaji, Magnus Johansson, Maria Tenje

## Abstract

Electroporation of dye-labelled bio-molecules has proven to be a valuable alternative to fluorescent protein fusion for single-molecule tracking in living cells. However, control over cell viability, electroporation efficiency and environment conditions before, during and after electroporation is difficult to achieve in bulk experiments. Here, we present a microfluidic platform capable of single-cell electroporation with *in situ* microscopy and demonstrate delivery of DNA into bacteria. Via real time observation of the electroporation process, we find that the effect of electrophoresis plays an important role when performing electroporation in a miniaturized platform and show that its undesired action can be balanced by using bipolar electrical pulses. We suggest that a low temperature of the sample during electroporation is important for cell viability due to temperature-dependant viscoelastic properties of the cell membrane. We further found that the presence of low conductive liquid between cells and the electrodes leads to a voltage divider effect which strongly influences the success of on-chip electroporation. Finally, we conclude that electroporation is intrinsically a highly stochastic process that is difficult to fully control via external parameters and envision that the microfluidic system presented here, capable of single-cell read-out, can be used for further fundamental studies to increase our understanding of the electroporation process.

## 1. Introduction

### 1.1. Modern biology sets new requirements for electroporation

Electroporation, or electropermeabilization, is the process in which electrical pulses are used to form pores in cell membranes to let otherwise impermeant biomolecules enter into the cell cytosol. The process was theoretically described in the early 1970s for isolated biomolecular lipid membranes (Crowley 1973) and in 1982, the method was practically demonstrated for the delivery of genetic material into mammalian cells (Neumann et al. 1982), and in the late 1980s, it was exploited for the delivery of plasmid DNA into bacterial cells (Dower, Miller, and Ragsdale 1988; Harlander 1987). Nowadays, electroporation of bacteria for DNA uptake, *i.e.* electrotransformation, is routine, and commercially available electroporation instruments are standard equipment in molecular biology and microbiology laboratories.

With more recent revolutions in fluorescence-based super-resolved microscopy, imaging of sub-cellular physiology and quantitative biochemical studies of molecular processes directly inside living cells have become possible (Q. Huang et al. 2021; Zalejski, Sun, and Sharma 2023). For single-molecule tracking, electroporation of biomolecules that have been fluorescently labeled with small organic dyes (Crawford et al. 2013; Volkov et al. 2018) has proven a valuable alternative to genetically fused fluorescent proteins. Compared to genetic transformation, in which electroporation is followed by a selection step to identify the successfully transformed cells, *in situ* electroporation for single-molecule fluorescence studies requires a more delicate balance between the survival rate and internalization efficiency as such a selection step is not available. This means that the technical advancement in microscopy, with corresponding paradigm shifts in life science research aiming for single-cell studies with single-molecule resolution, puts new demands on cell electroporation.

At the same time, significant progress has been made in micro- and nanostructure fabrication, allowing us to manufacture systems to easily study and follow cells and their progeny in a controlled environment (Anggraini et al. 2022). Microfluidic devices may, in addition, provide single-cell control, direct *in situ* observation of the electroporation process under the microscope, and potential integration of electroporation into lab-on-a-chip multifunction platforms with high throughput and reduced reagent consumption. Hence, microfluidics holds the potential to enable single-cell transformation and/or internalization of fluorescently labeled biomolecules through electroporation, thereby combining the achievements of these two independent fields.

### 1.2. Physical principles of electroporation

To understand potentially important aspects for building a miniaturized electroporation platform, we must first consider the basic principles behind the electroporation phenomenon. We have briefly summarized the process below. For more detailed reviews on electroporation, the interested reader is referred to (Kotnik et al. 2019; Teissie 2014).

All living cells possess a resting membrane potential defined as an electrical potential difference between external and internal membrane surfaces (typically −40 to −70 mV for eukaryotic cells (Kotnik et al. 2019) and around −140 mV for the bacterium *E. coli* (Lo et al. 2007)), making it possible for cells to perform chemical and mechanical work. In a simplified physical model, the cell represents a compartment made of a low-conductivity layer (membrane) filled with an electrolyte (cytoplasm). The conductivity of the cell membrane is several orders of magnitude lower than that of the cytoplasm and the surrounding media. This means that if a cell is exposed to a homogeneous external electric field, the electric field is concentrated within the membrane, resulting in an induction of transmembrane voltage (TMV) in addition to the existing resting potential (Kotnik and Pucihar 2010). In the case of a simple cell geometry, and assuming steady-state conditions (*i.e.* that the electrical field is applied longer than the characteristic charging time of the membrane), the TMV can be expressed analytically by solving Laplace’s equation for an electrostatic potential with realistic boundary conditions (Kotnik and Pucihar 2010). For an ellipsoidal cell, the solution is given by Schwan’s equation (Bernhardt and Pauly 1973; Kotnik and Pucihar 2010):

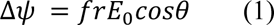

Where *f* is a form-factor (1.5 for spherical cell), *r* is the cell radius along the field, *E_o_* is the applied external field, and *θ* is the polar angle measured from the center of the cell with respect to the direction of the field.

When the TMV is high enough (typically a few hundred mV (Towhidi et al. 2008)), detectable pores begin to appear. As the rate of the underlying physical processes leading to electroporation (*e.g.* lipid rearrangements) increases nonlinearly with the field amplitude, there is, strictly speaking, no sharp threshold for the TMV to create pores and only an approximate value of TMV can be given (Kotnik et al. 2019). Pores become detectable on the nanosecond to microsecond time scale (depending on the experimental conditions and detection method) and expand while the pulse lasts (Frey et al. 2006; Hibino, Itoh, and Kinosita 1993; Pucihar et al. 2008). The formed pores can either self-seal or remain irreversibly open. In the case of irreversible pores, the integrity of the cellular membrane is broken down, the cell dies, and the cellular content is released into the environment. Such treatment is used for extraction of cellular components, medical treatments (*e.g.* killing cancer cells), and sterilization in the food industry (Haberl et al. 2013). If the cell should remain alive following electroporation, self-healing pores are required. These types of pores are much more difficult to generate due to their metastability. Partial recovery of reversible pores takes place on the microsecond to millisecond time scale (Hibino, Itoh, and Kinosita 1993; Prausnitz et al. 1995; Pucihar et al. 2008), making it a much slower process than pore formation. Full re-sealing does not occur until seconds or hours after the pulse has been removed, depending on environmental factors such as temperature (Lopez, Rols, and Teissie 1988; Pucihar et al. 2008; Shirakashi et al. 2004). Besides TMV induction, the external electrical field induces electrophoresis of charged molecules through the pores, resulting in intracellular delivery of otherwise impermeant biomolecules (*e.g.* DNA and proteins). The latter is widely exploited for delivery of DNA for genomic modifications, but also more recently for delivery of fluorescently labeled biomolecules for single-molecule tracking studies in live cells (Plochowietz et al. 2017; Rivera et al. 2021; Volkov et al. 2018, 2022).

In addition to the external electrical field, many other factors such as temperature and membrane composition influence pore formation and dynamics. Besides physical alteration of the lipid bilayer, pore formation can also induce additional damage to the cell, such as oxidative damage of the phospholipids (Breton and Mir 2018), and conformational changes or denaturation of membrane proteins (Chen 2006). The physiological stage of the cells has also been shown to greatly affect cell viability after the pulse (Rols et al. 1998). Thus, the electroporation process is complex and far from fully understood, and is therefore experimentally challenging in terms of efficiency and cell viability and requires increased experimental attention.

### 1.3. Electroporation of bacteria

As a rule, the electrical and environmental parameters for electroporation are adjusted for each cell type and desired application. Electroporation of eukaryotic cells for DNA delivery is typically performed in a salt-containing electroporation buffer with a field strength of 0.01-0.1 kV/mm (Kumar, Nagarajan, and Uchil 2019; Potter and Heller 2003). Interestingly, DNA delivery in bacteria generally requires about an order of magnitude higher field strength (∼1 kV/mm for *E.coli* (Dower, Miller, and Ragsdale 1988)). This is partially due to the smaller cell size, resulting in less efficient build-up of TMV under the same external field (Equation 1) (Choi, Khoo, and Hur 2022), but also due to the different composition of the cell membranes. Bacteria, such as *E. coli*, contain two lipid bilayers with a peptidoglycan cell wall in between (Silhavy, Kahne, and Walker 2010), whereas mammalian cells only have a single lipid bilayer and no cell wall (Fowler, Roush, and Wise 2013). Due to the much higher electrical fields required for bacteria, electroporation is typically performed in deionized water or in 10 % glycerol (cryoprotectant for deep-freeze storage) to prevent high current and Joule heating of the sample (Chang et al. 1991).

When an electrical field is applied, it induces a dipole on rod-shaped cells that experience a torque and orient along the field lines, with a typical half-time shorter than ∼1 ms (N. Eynard et al. 1998; Nathalie Eynard et al. 1992). The orientation of the cell with the electrical field, in turn, changes the amplitude of the induced TMV due to a change in the form-factor *f* and the equivalent radius *r* (Equation 1). For example, for an ellipsoidal cell resembling *E. coli* (axial ratio 1:1:3), the TMV induced in the region facing the electrode (*θ = 0*) increases twice upon cell rotation from perpendicular orientation (*f* = 1.8, *r* = 1) to one co-aligned (*f* = 1.2, *r* = 3) with the electrical field (N. Eynard et al. 1998). Thus, alignment of cells along the field is more favorable for electroporation of rod-shaped bacteria at a given voltage applied to the electroporation electrodes.

### 1.4. Miniaturization of electroporation devices

Cell electroporation is routinely performed in electroporation cuvettes with a 1‒4 mm gap between electrodes. Miniaturization of the electroporation device may provide better environmental, spatial, and temporal control over the process, thus improving the efficiency (Choi, Khoo, and Hur 2022; Movahed and Li 2011). A smaller distance between the electrodes also means that the field strength required for cell permeabilization can be achieved at lower voltages. A lower voltage, in turn, leads to decreased Joule heating, which is also beneficial when handling biological samples, and it further reduces the rates of undesired electrochemical reactions by the electrodes (*e.g.* electrolysis of water and reactions of dissolved components with the electrode material) (Choi, Khoo, and Hur 2022; Movahed and Li 2011).

Although there have been a wide number of microfluidic platforms developed for single-cell electroporation of eukaryotic cells (for review see (Kar et al. 2018) and references therein), miniaturized devices for bacterial electroporation are rarely reported. Further, most of them use irreversible electroporation for bacterial inactivation (Fox et al. 2005; Pudasaini et al. 2021) or extraction of cell content (Bao and Lu 2008; Blank et al. 2019; Ma et al. 2016; Poudineh et al. 2014; H. Y. Wang, Bhunia, and Lu 2006), whereas some reports focus on reversible electroporation of bacteria for delivery of plasmid DNA (Garcia et al. 2017; P. H. Huang et al. 2022; Madison et al. 2017; Shih et al. 2015). However, the distances between electrodes in all these devices are tens to hundreds of micrometers, thus requiring a high voltage, and all of the devices have a flow-through design which does not allow for single-cell treatment and *in situ* analysis.

To understand the electroporation process better and approach controllable *in situ* single-cell electroporation of bacteria for super-resolution microscopy, we set out to systematically investigate the external parameters affecting electroporation success. For this purpose, we developed a microfluidic platform in which the cell orientation in the applied electrical field could be controlled, and where different field strengths could be applied. The system includes temperature and liquid exchange control and is compatible with microscopy for *in situ* observation of the electroporation process, and consecutive single-molecule fluorescence microscopy of the electroporated cells. Additionally, we compared the on-chip electroporation results with those obtained from in-bulk electroporation under analogous conditions.

## 2. Results and discussion

### 2.1. Microfluidic platform for *in situ* electroporation

In the design of the miniaturized electroporation device and protocol, we followed the common procedure for bulk electroporation of *E. coli* cells in electroporation cuvettes (Chang et al. 1991). We used a PDMS-based microfluidic chip, also known as the “mother machine” (P. Wang et al. 2010), to trap bacteria in tight microfluidic channels measuring 1.2 *x* 1.2 *x* 50 μm^3^ in size. The channels are constricted at one end, preventing the cells from escaping while still allowing a flow of growth medium and reagents over the cells (Figure 1 and S1). A similar microfluidic device has been used previously by us and others to grow and observe bacterial cells, for example when exposed to antibiotics (Baltekin et al. 2017; Rivera et al. 2021). Compared to the previous mother machine chip, our new device was modified in two ways. First, we used integrated liquid cooling channels (Khaji and Tenje 2022) to cool down cells and liquid before and during the electrical pulse. Second, we structured thin-film platinum electrodes on the glass surface at the bottom of the microfluidic chip. The electrodes were carefully aligned perpendicularly to the cell traps. This way, the cells were aligned with the electrical field already before the pulse (compared to bulk electroporation where rod-shaped bacteria are in random orientation in solution before the pulse and become polarized and oriented along the field during the first milliseconds of the pulse (N. Eynard et al. 1998; Nathalie Eynard et al. 1992)). Cells tightly loaded in the cell traps in a pole-to-pole fashion thus represent a one-dimensional strand orientated parallel to the field. Based on calculations for spherical cells packed similarly, where the induced TMV is predicted to be 1.5 times lower than the TMV induced in isolated single cells under the same conditions (Susil, Šemrov, and Miklavčič 1998), we expected that the chosen geometry of our microfluidic device would result in a similar decrease in the induced TMV compared to isolated single cells, which can be compensated with a slight increase of applied voltage.

**Figure 1.**
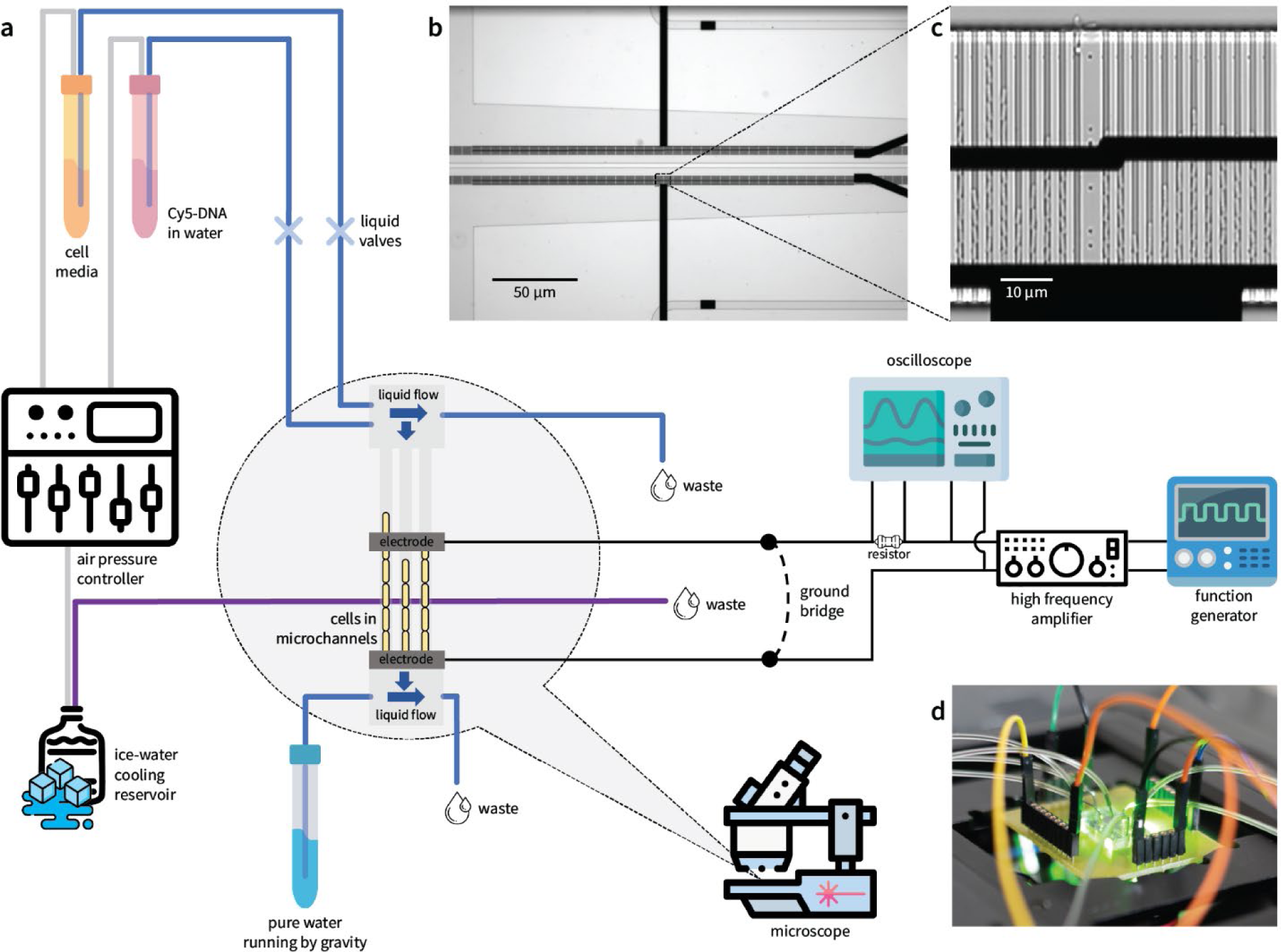
Experimental setup for *in situ* study of bacteria electroporation. (a) Schematic overview of the setup. (b) Microphotograph of the PDMS “mother machine” chip with cell traps crossed by two blocks of pulsing electrodes: one with 20 µm distance between electrodes (upper) and one with stepwise variable distances of 16, 18, 20, and 22 µm between the two electrodes (lower). (c) Cell traps with loaded *E. coli* cells crossed by an electrode block with stepwise variable distance of 18 and 20 μm. (d) Photograph of microchip installed on the microscope stage with connected liquid tubing and electrical wires. The figure includes images from flaticon.com.

The thin-film electrodes were made of platinum and a thin layer of tantalum was used for adhesion. The electrodes were patterned on the glass bottom of the microfluidic chip via a lift-off process including UV lithography and metal sputtering. The chip holds four independent blocks of electrodes with either 20 µm between the pulsing electrodes, or with stepwise variable distances of 16, 18, 20, and 22 µm (Figure 1 b, c). The blocks with variable distances between the electrodes allowed us to pulse cells with four different field strengths simultaneously. To prevent any short-circuiting of the system and ensure that practically all the voltage drop occurred between the electrodes across the loaded cells, care was taken to ensure that the highly conductive cell growth media was fully replaced by low conductive liquid before the electrical pulse was applied. As a measure of caution, all electrodes on the chip were kept grounded at all times, except the pulsing electrode block for which the ground bridge was manually removed a few seconds before the pulse.

Devices for routine bulk electroporation of bacteria supply an exponentially decaying pulse with a time constant dependent on the sample’s conductivity (Chang et al. 1991). In our set-up, we aimed for better control over the pulse shape and duration, and for this reason, we chose a square-shaped pulse. A ms-long square electrical pulse was produced using a function generator followed by amplification using a high-frequency amplifier. This produced a pulse of 20-30 V between the microelectrodes, creating a field strength of ∼1-1.5 V/µm, which is similar to the field strength typical for bulk procedures (1 V/µm = 1 kV/mm). A two-channel oscilloscope was connected to monitor pulse shape, and passage of the electrical current through the circuit measured as a drop in voltage on a resistor in series with the electroporation circuit (Figure 1).

As a model system, we chose to electroporate a common *E. coli* laboratory cell strain (MG1655) and internalize 14-mer DNA oligonucleotides labeled with the fluorescent Cy5 dye (Cy5-DNA) for observation under a fluorescence microscope. To mimic the typical bulk electroporation procedure (Chang et al. 1991), cells were grown at 24° C until they reached a stable exponential growth phase and then loaded into the microchip. Immediately after cell loading, the on-chip cooling system was activated, cells were cooled down to 6° C, and cell media was changed to deionized water containing Cy5-DNA (Supplementary Video 1). Preliminary experiments, with the addition of 10 % glycerol, which is widely used in bulk electroporation (Chang et al. 1991), resulted in problems with liquid exchange of the microchannels due to the solution’s high viscosity. Hence, glycerol was omitted in all subsequent on-chip experiments. After the electrical pulse was applied to one of the electrode blocks, the water/Cy5-DNA solution was changed to cell media, the cooling system was deactivated, and cells were imaged for detection of internalized Cy5-DNA and cell growth. The flow of liquid supplied to the cells was managed by a pressure-control unit and remotely controlled valves for rapid switching of liquid before and after the pulse.

As a first test, we explored the effect of a 1 ms square monopolar pulse (a typical pulse trace is shown in Figure 4c) with a field strength of 1-1.4 V/µm in our miniaturized platform. During the experiment, the microfluidic chip was placed under a laser scanning confocal fluorescence microscope, which allowed us to follow cell loading, exchange of liquid, the electroporation process, and finally, the results of electroporation in terms of Cy5-DNA internalization and cell viability. The Cy5 fluorescence signal was detected and bright-field images were captured to record growth every 5 min for 60 min after the pulse. Fluorescence images were processed to construct a fluorescence binary mask, which was then overlaid with the corresponding bright field image, indicating in which cells Cy5-DNA was internalized. We did not observe Cy5 signal in the cells before electroporation (Figure 2a and S2a), nor in any cells outside the active electrodes or between the grounded electrodes after the active electrodes were pulsed (negative control, Figure S2b). Moreover, almost all cells in the negative control (> 96 %) grew normally after being subjected to the media and temperature shifts of the electroporation procedure (Supplementary Video 2). On the other hand, among the cells located between the pulsed electrodes, and thus subjected to the electrical field, some exhibited a visible Cy5 signal, and some were growth impaired (Figure 2b, 2d, 2e, and Supplementary Video 3).

**Figure 2.**
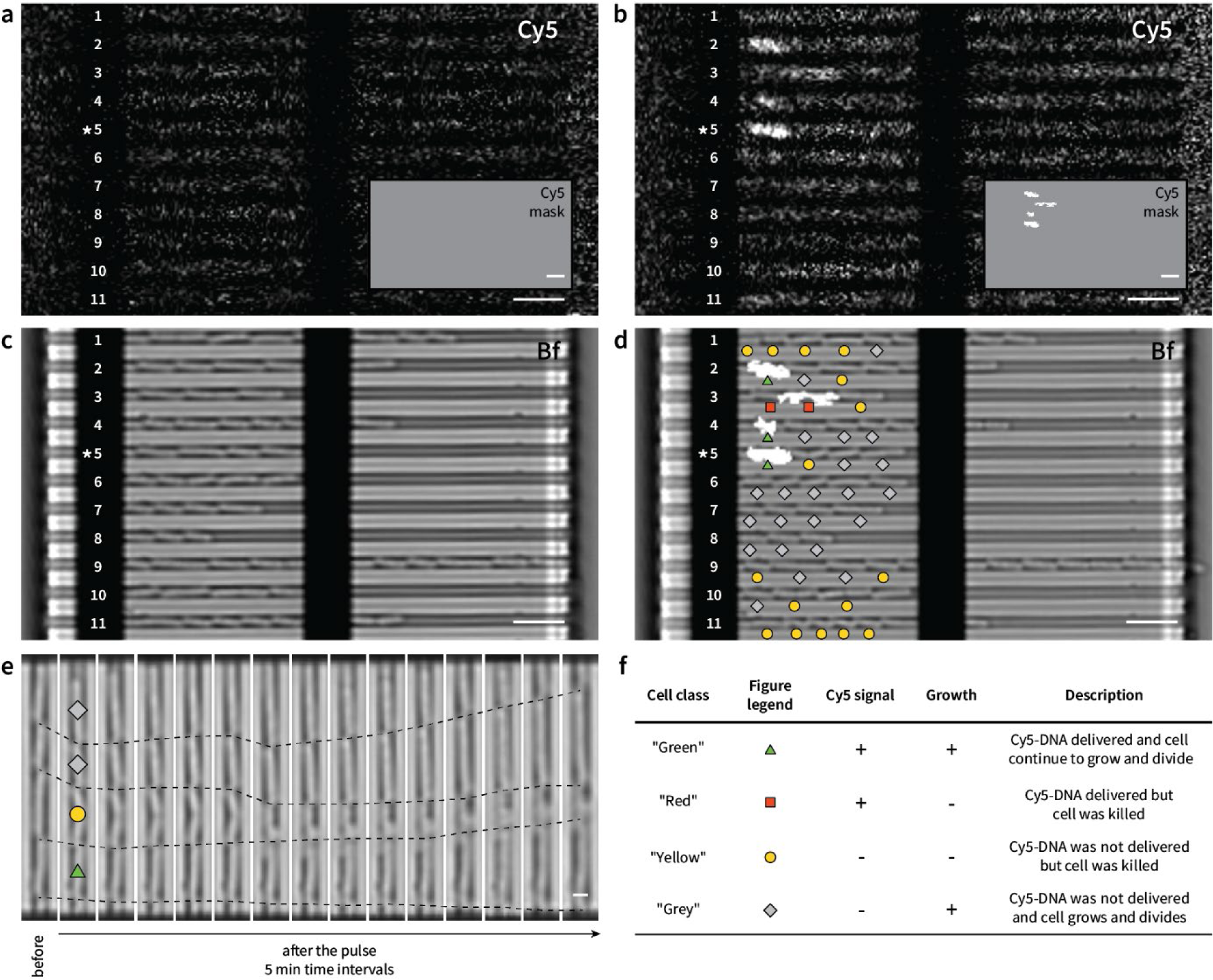
Microphotograph of the microchip with loaded cells before (a, c) and after (b, d) electroporation with a 1 ms 1.2 V/μm monopolar pulse. Cy5-fluorescence (a, b), Cy5-fluorescence mask (inset in (a, b)) and bright field images, overlaid with white Cy5-fluorescence mask (c, d). Cell traps are numbered from 1 to 11. After electroporation, cells were classified as marked with the different symbols in (d) and (e) based on detectable Cy5 signal and cell growth, as explained in (f). (e) Cells loaded in trap number *5 imaged before electroporation and after electroporation with 5 min intervals. Cell poles in sequential frames are connected with dashed lines as a guide to the eye. Scale bars (in white) are 5 μm in (a-d) and 1 μm in (e). Cell growth time-lapse corresponding to (d) is shown in Supplementary Video 3.

The pulsed cells were sorted into four classes based on the presence or absence of detectable Cy5 signal and observed growth pattern (Figure 2f). In our nomenclature, both “green” and “red” cells had internalized Cy5-DNA and the “green” cells continued to grow whereas the “red” cells did not. “Yellow” and “grey” cells did not have any internalized Cy5-DNA, and the “grey” cells continued to grow, whereas the “yellow” cells did not. Under these pulse conditions, we were able to find cells with internalized Cy5-DNA, which also resumed growth (*i.e.* “green” cells, Figure 2e).

### 2.2. On-chip electroporation with monopolar pulse

#### 2.2.1. Number of cells loaded into the traps affect electroporation efficiency

From the fluorescence images obtained after electroporation, we immediately noted that the result of the process was highly heterogeneous; we could find cells of different classes in different traps and even within the same cell trap (Figure 2d), even though all cells had the same orientation and experienced nearly identical conditions during the pulse. We first hypothesized that the difference in the number of cells loaded in the traps could influence the electroporation result. To investigate this hypothesis, we classified cell traps based on the cell loading pattern and analyzed cells from each class separately. We selected three types of loading patterns, which we for simplicity call “underfilled”, “normal”, and “overfilled” depending on the length *C* occupied by cells with respect to the electrode-to-electrode distance *D* and total trap length *L,* as explained in Figure 3a.

**Figure 3.**
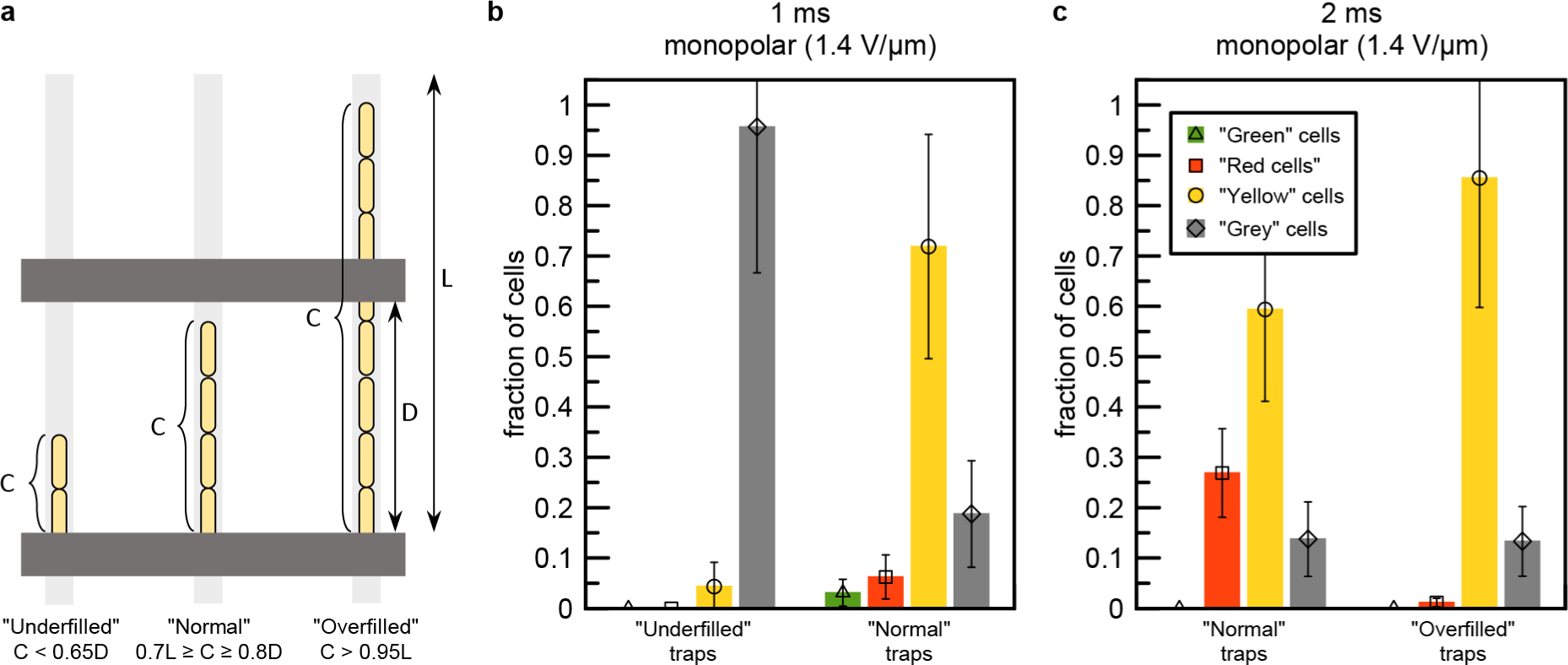
Number of cells loaded into the traps affect cell survival and permeabilization efficiency. (a) Three types of cell traps were selected for analysis based on the number of loaded cells with respect to the distance between electrodes *D* and total channel length *L* (from trap entrance to constriction). (b) Electroporation results for cells in “underfilled” and “normal” cell traps electroporated with 1 ms 1.4 V/μm monopolar pulse. (c) Electroporation results for cells in “normal” and “overfilled” cell traps electroporated with 2 ms 1.4 V/μm monopolar pulse. Cell classification are explained in Figure 2f. Error bars represent combined extrapolated reproducibility and measurement errors (see Experimental section).

We first considered cells from “underfilled” and “normal” traps, disregarding the “overfilled” traps. Analysis of the Cy5 signal and cell survival revealed a striking difference between these two subsets of cells. For example, after a 1.4 V/µm 1 ms pulse, more than 95 % of the cells from the “underfilled” traps were unaffected (“grey” cells, Figure 3b), whereas in “normal” traps, more than 75 % of the cells were killed (“yellow” and “red” cells, Figure 3b). We hypothesize that this difference is due to the different voltages the cells in “underfilled” and “normal” traps are exposed to. Since the conductivity of the liquid in the microchannel (*i.e.* deionized water containing DNA), around 0.2 mS/m (Supplementary Note 1), is much lower than the conductivity of a cell (30 mS/m (Carstensen et al. 1965)), a voltage divider effect occurs. That is, the water in the partially filled traps results in a significant voltage drop, and 12 the remaining voltage is not enough to produce a detectable effect on the cells. We observed a similar effect with microfluidic devices in which the electrode alignments were not perfect, and hence, one of the electrodes was at a greater distance from the cell traps in the back channel. This created a region filled with low-conductive liquid between the cells and the electrode (Figure S3). In this case, the effect of the pulse on cells was very mild or non-existent, thus confirming that the difference between “underfilled” and “normal” traps is governed by a voltage divider effect rather than mechanical or fluidic effects. A similar voltage divider effect, inhibiting electroporation, has been observed for electroporation electrodes coated with a thin dielectric film (Wassermann et al. 2016).

To further investigate the effect of cell trap loading, we extended the cell-loading time to trap a higher number of cells in each microchannel. In an attempt to improve internalization of Cy5-DNA in the cells, the pulse time was also increased to 2 ms while the field strength remained at 1.4 V/µm. By comparing results from “normal” traps for 1 ms and 2 ms pulses (Figure 3b and c), we found that a longer pulse time resulted in a much higher fraction of cells with internalized Cy5-DNA (∼27 % *vs.* ∼9 % at 1 ms), even though nearly all cells containing Cy5-DNA were dead (“red” cells Figure 3c). However, we noticed that cells in “overfilled” traps did not internalize Cy5-DNA at all (∼1 % of “red” cells and 0 % of “green” cells), despite the rather harsh pulse conditions, and most of the cells also died (Figure 3c). We speculate that the densely packed cells in the overfilled channels physically obstruct Cy5-DNA from entering the microchannel before or during the electroporation pulse. However, we did not quantify this effect further in our experiments.

To minimize the effect of cell loading on electroporation, we sorted all data for further analyses based on loading type (Figure 3a) and retained only data from “normal” traps where the voltage divider effect was less significant and Cy5-DNA internalization uninhibited. Nevertheless, the results of electroporation were highly heterogeneous between cells in the same trap and between cells in neighboring traps filled to a similar extent (see for example traps number 5, 6, and 10 in Figure 2d).

#### 2.2.2. Higher field strength and longer pulse duration increase permeabilization efficiency

When varying the field strength of a 1 ms pulse from 1.0 to 1.4 V/µm by exploiting the electrodes with stepwise variable distances, we could observe a clear trend in electroporation outcome. As expected, the fraction of “grey” cells (unaffected) decreased with increasing field strength (Figure 4a). Under these conditions, the fraction of “green” cells was only ∼3-5 %, while the fraction of “yellow” cells was ∼15-60 %, depending on the field strength (Figure 4a). We attempted to increase the yield of “green” cells by increasing the pulse duration, from 1 ms to 2 ms, but it resulted in a smaller fraction of “grey” cells instead without any significant change in the fraction of “green” cells (Figure 4b). By comparing 1 ms and 2 ms pulse results, we see that the outcome of electroporation as a function of field strength follows a similar trend (Figure 4a, b). However, with a longer pulse, the same effects were achieved at a lower field strength. A similar compensatory relationship between pulse duration and field strength has previously been observed for bulk electroporation (Dower, Miller, and Ragsdale 1988).

**Figure 4.**
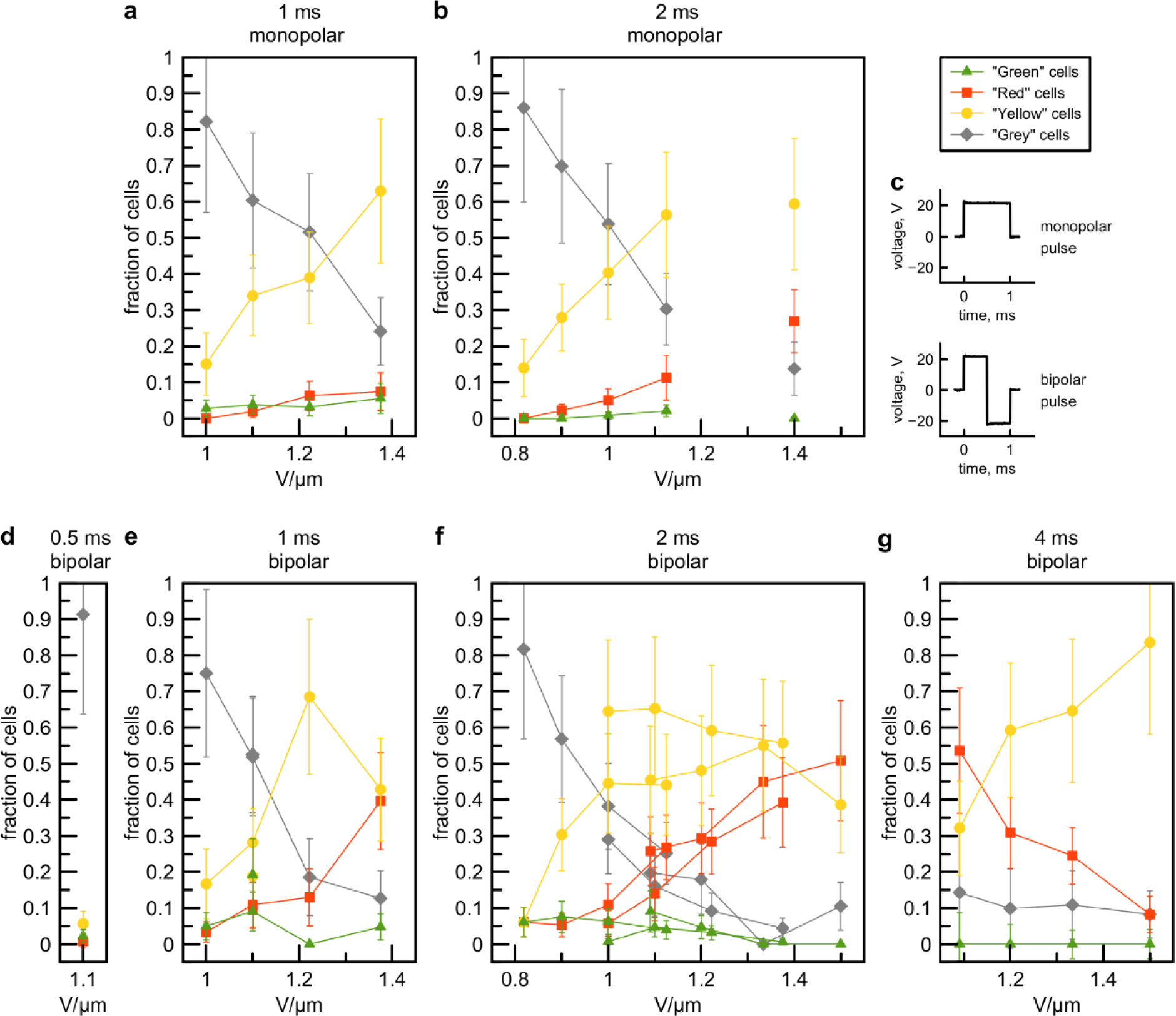
Results of electroporation obtained with monopolar (a, b) and bipolar (d-g) square pulses with varied field strength and pulse duration. Cell classification is described in Figure 2f. Straight lines as a guide to the eye connect experimental points obtained in one experiment from a block of electrodes with stepwise variable distances. (c) Typical traces of monopolar (upper) and bipolar (lower) 1 ms square pulses of 22 V amplitude applied to electroporating electrodes as registered by the oscilloscope. Error bars represent combined extrapolated reproducibility and measurement errors (see Experimental section).

#### 2.2.3. Electrophoresis must be considered, for both cells and molecules

With the microfluidic platform installed under the microscope, we had a unique possibility to visually observe the electroporation process *in situ*. For this purpose, we imaged the microfluidic system with bright-field and fluorescence illumination at 19 ms/frame, when cells experienced a 2 ms electrical pulse (Figure 5a and Supplementary Video 4). Even though the time resolution of the acquired data was lower than the actual pulse time, the effect of the pulse was clearly visible. Before and after the pulse (*i.e.* frames 1 and 3, respectively), the cells were located at the end of the cell trap due to the constant liquid flow. However, during the pulse (frame 2), the cells moved significantly towards the trap entrance, before moving back again, colliding with the cell trap constriction and each other. We attribute this movement to electrophoresis of the cells in the electrical field created between the electrodes. *E. coli* cells in low ionic liquid are negatively charged (Güler and Oruç 2021) and thus move towards the positive electrode. This electrophoretic movement could be reversed by reversing the polarity of the electrodes (Figure S4).

**Figure 5.**
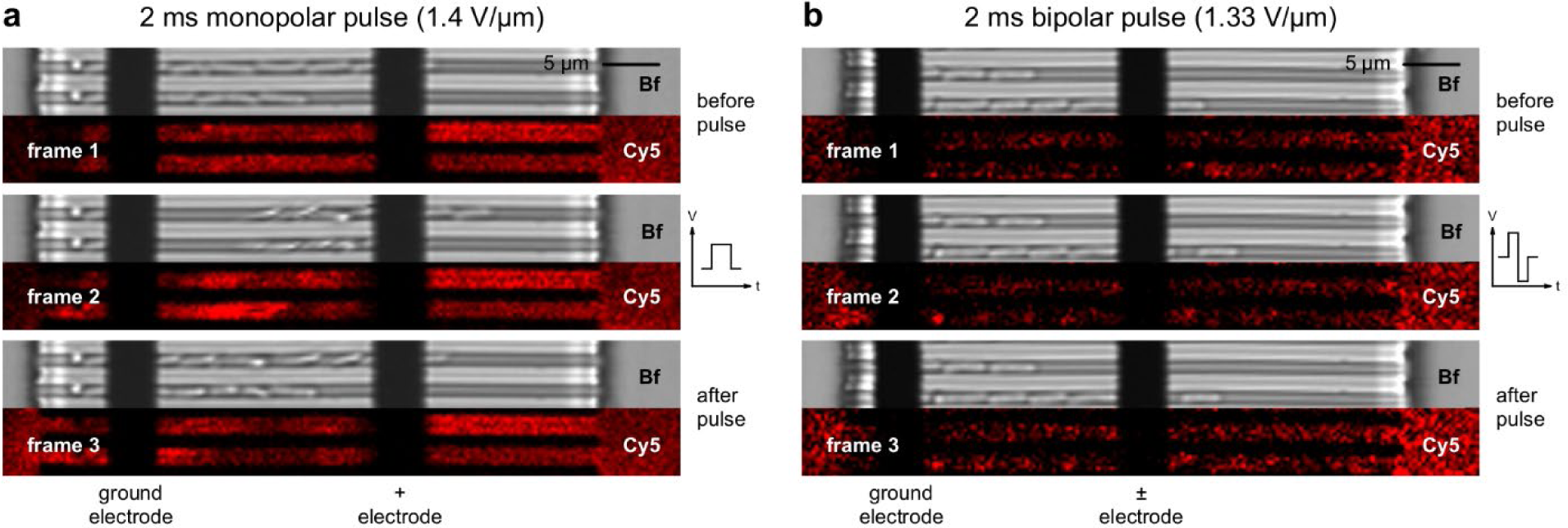
*In situ* observation of electroporation with 2 ms square electrical pulse recorded at 19 ms per frame with bright field and Cy5-fluorescence imaging. Ground and pulsed electrodes are marked. (a) Monopolar pulse of 1.4 V/μm. (b) Bipolar pulse of 1.33 V/μm. Corresponding time-lapse videos are shown in Supplementary Video 4 and 5.

Although we could not obtain an instant picture of Cy5-DNA electrophoretic movement between the electrodes with the time resolution of our optical setup, probably because this process is much faster than the movement of the cells due to the higher charge density and smaller size of the 14-mer DNA, we did observe local changes in Cy5 signal density during and after the pulse (Figure 5a and Supplementary Video 4).

We thus speculate that pulse-induced electrophoresis decreases the local Cy5-DNA concentration between active electrodes, as well as misplaces cells from the optimal position between them. Both these effects could potentially decrease the efficiency of Cy5-DNA internalization and lead to a lower fraction of live cells with internalized Cy5-DNA (“green” cells), and should therefore be minimized.

#### 2.2.4. Bipolar pulses increase the electroporation efficiency

In an attempt to increase the yield of live cells with internalized Cy5-DNA (“green” cells) by reducing the counteractive effects of electrophoresis, we next investigated the electroporation outcome of a bipolar pulse (a typical trace is shown in Figure 4c). We hypothesized that a quick switch of polarity, as provided by the bipolar pulse, would counteract the potentially negative effect of electrophoresis during the monopolar pulse. Indeed, when a bipolar square pulse with similar field strength and the same total duration as the previous monopolar pulse was applied, the cells did not move as much during and after the bipolar pulse (Figure 5b and Supplementary Video 5). We also observed a more homogenous Cy5-DNA concentration in the microfluidic channels.

We noticed that a 1 ms bipolar pulse resulted in a larger total fraction of cells with internalized Cy5-DNA (sum of “green” and “red” cells) compared to a monopolar pulse of the same amplitude and duration, with the difference generally more pronounced as field strength increased (e.g. ∼45 % of cells with electroporated Cy5-DNA for bipolar, compared to ∼13 % for monopolar, at 1.4 V/µm, Figure 6a). Consequently, the fraction of electroporated and live (“green”) cells was higher with a bipolar pulse (10-20 % in 2 repetitions) compared to the result with a monopolar pulse (∼4 % in 1 experiment, Figure 4a, e).

**Figure 6.**
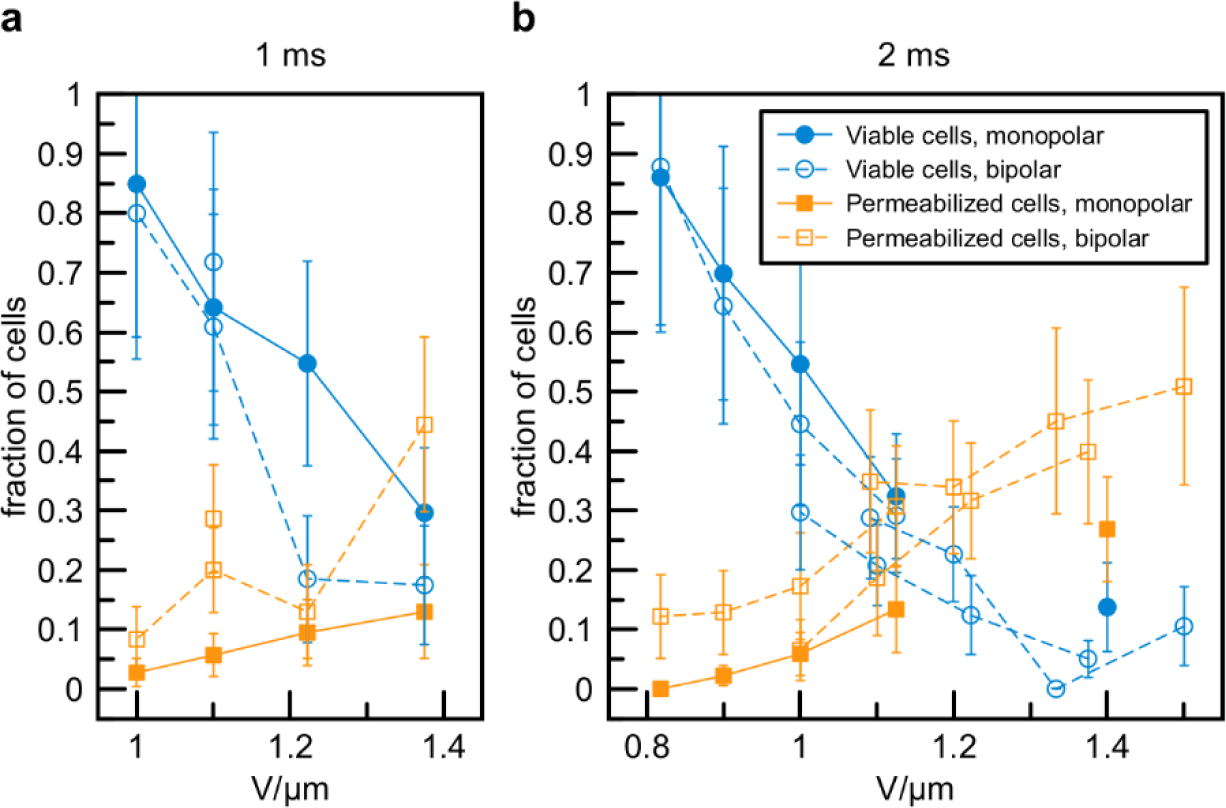
Results of electroporation obtained with 1 ms (a) and 2 ms (b) square monopolar and bipolar pulses. Cells are classified as viable based on growth after the pulse (sum of “green” and “grey” classes according to Figure 2f) and as permeabilized based on Cy5 signal (sum of “green” and “red” classes). Straight lines as a guide to the eye connect experimental points obtained in one experiment from a block of electrodes with stepwise variable distances. Error bars represent combined extrapolated reproducibility and measurement errors (see Experimental section).

To further elaborate the difference between monopolar and bipolar pulses and validate reproducibility of our data we performed a few experiments with longer, 2 ms bipolar pulses of different amplitude (18, 22, 24 V) applied to electrodes with stepwise variable distance. This allowed us to cover the range of 0.8 - 1.5 V/µm field strength with partially overlapping conditions between experiments. Similarly to 1 ms pulses, 2 ms bipolar pulses resulted in more efficient internalization of Cy5-DNA compared to monopolar pulses (mean 16 % for bipolar *v.s.* mean 5 % for monopolar over the range of 0.8 - 1.1 V/µm) (Figure 6b). Increased internalization efficiency also led to increased fraction of “green” cells (mean 5 % for bipolar *v.s.* mean 1 % for monopolar over the range of 0.8 - 1.1 V/µm (Figure 4b, f).

Due to the complexity in fabrication and operation of the microfluidic chips, the number of conducted experiments is unfortunately limited in this study. Consequently, certain experimental conditions have not been attempted to reproduce. However, the reproducibility error can be assessed and extrapolated to all experiments from the data where replicas exist. To quantify the reproducibility errors, we computed the coefficient of variation (ratio of standard deviation to the mean) for data points obtained under close field strengths (1.091, 1.1 and 1.125 V/µm) at 2 ms bipolar pulses in three independent experiments (Figure S5). We found that the typical coefficient of variation does not exceed 0.3 for values > 0.2 and become ∼0.5 for smaller values. (*i.e.* the relative error due to experiment-to-experiment variability is 30-50%). We also combined this error with the measurement error caused by limited sample size for individual on-chip experiments (see Experimental section). Though we note that some datapoints for monopolar and bipolar pulse results are closely located relative to the error margins, we see reproducible trends for two different pulse durations (1 and 2 ms, Figure 6) and in both cases, bipolar pulses resulted in slightly better Cy5-DNA internalization.

We further varied the duration of the bipolar pulse and found that a pulse of 0.5 ms, at 1.1 V/µm, was too brief to produce a significant effect on the cells (Figure 4d). More than 90 % of the cells recovered after the pulse without internalizing Cy5-DNA (“grey” cells). On the other hand, a 4 ms pulse was too long to produce any detectable fraction of “green” cells (Figure 4g). Interestingly, an increase of the field strength with 4 ms bipolar pulses inhibited Cy5-DNA delivery, *i.e.* the fraction of “red” cells decreased, whereas the opposite was observed with shorter pulses.

Finally, by instead plotting the outcome of electroporation with bipolar pulses at 1.1 V/µm field strength (the field strength that yields the maximum number of “green” cells) as a function of pulse duration, we saw a clear trend of a monotonically decreasing fraction of “grey” cells and an increasing fraction of “red” cells with longer pulses (Figure 7). The fractions of “yellow” and “green” cells, on the other hand, appear to have maxima at around 2 ms and 1 ms respectively (Figure 7).

**Figure 7.**
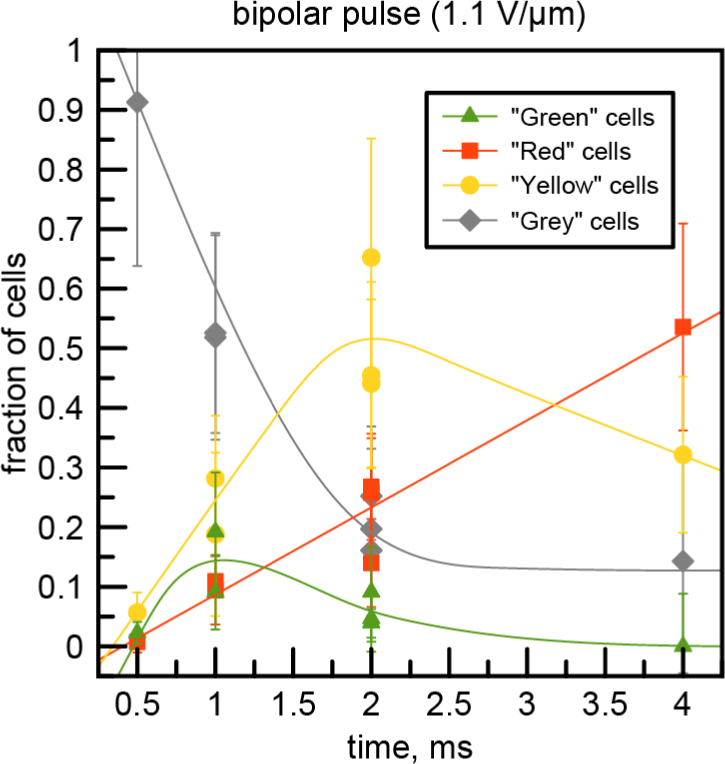
Results of electroporation with square bipolar pulse at 1.1 V/μm field strength as a function of pulse duration. Each data point represents an individual experiment. Cell classifications are described in Figure 2f. Solid lines are drawn as a guide to the eye. Error bars represent combined extrapolated reproducibility and measurement errors (see Experimental section).

#### 2.2.5. High cell viability requires low temperature during electroporation

To achieve high electroporation efficiency in bulk electroporation, it is generally recommended that cells are cooled down in ice, washed with cold water or 10 % glycerol, and electroporated in an ice-cold cuvette (Chang et al. 1991). As mentioned previously, our microfluidic setup has cooling capability, which was used in the experiments presented so far. Based on known pulse parameters, measured current, and the volume of liquid between electrodes in the microfluidic channels, we estimated that for a typical pulse of 1 ms, 22 V, and 20 µA, the temperature of the liquid should not increase to more than 5 °C (Supplementary Note 2). We note that this is an upper estimate since it does not account for energy dissipation during the pulse, nor heat transfer due to the constant liquid exchange on the chip. The estimated temperature increase is expected to be too small to affect cell viability, and hence, we were motivated to test whether cooling of the chip really was essential for the electroporation outcome. To investigate this, we electroporated cells with 1 ms bipolar pulse at 1-1.4 V/µm following the regular protocol, but on a microfluidic chip without the cooling system connected. The temperature of the chip was 28±2 °C during liquid exchange and pulsing. As in our previous experiments, the control cells located on the same chip, but not subjected to an electrical field, survived after liquid change, had no internalized Cy5-DNA, and grew normally. The cells between the electroporation electrodes, on the other hand, were nearly all killed after the pulse (> 95 % of “yellow” and “red” cells, Figure 8a), with Cy5-DNA delivered to a substantial fraction of them (“red” cells, ∼10-40 %, Figure 8a). This result clearly demonstrates that maintaining a low temperature during the pulse is essential for cell viability, although our calculations showed that the Joule heating of the entire volume between electrodes is insignificant.

**Figure 8.**
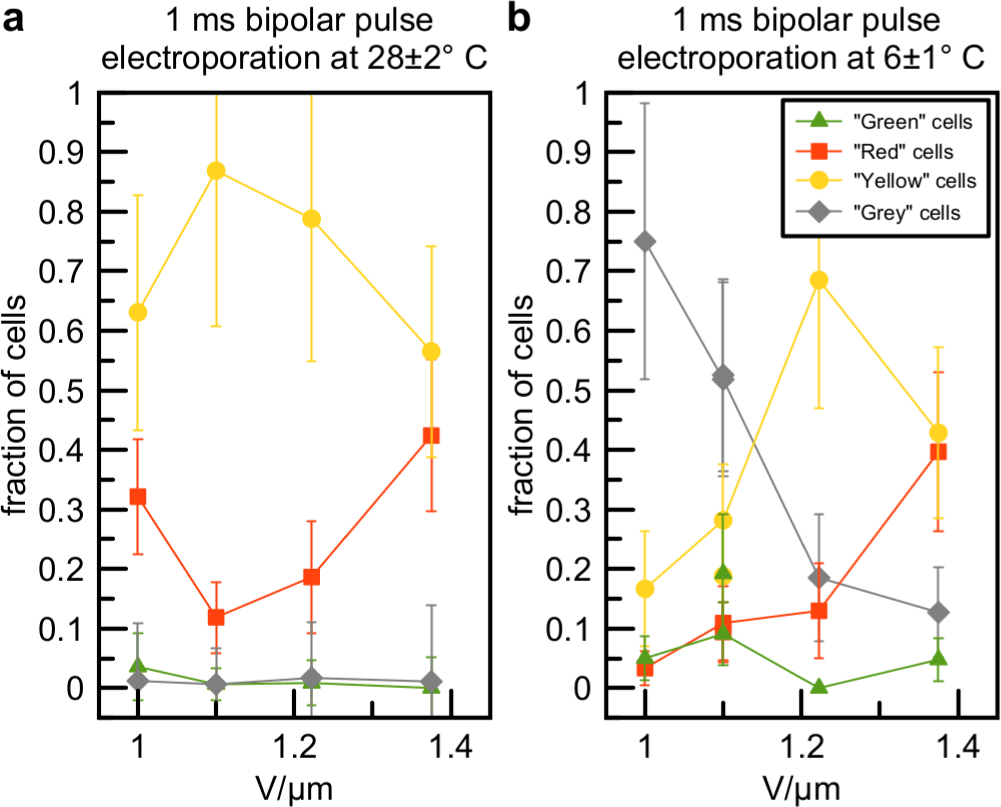
Results of electroporation with 1 ms square bipolar pulse at 28±2 °C (a) and 6±1 °C (b). Straight lines as a guide to the eye connect experimental points obtained in one experiment from a block of electrodes with stepwise variable distance. Cell classifications are described in Figure 2f. Data in (c) are the same as in Figure 4e and shown for comparison. Error bars represent combined extrapolated reproducibility and measurement errors (see Experimental section).

One possible explanation for the high fraction of dead cells could be the effect of local Joule heating caused by the electrical current through the opened electropores, which may cause local membrane protein denaturation and subsequent cell death (Tsong 1990). We also speculate that the observed lower cell viability could be caused by a higher number of irreversible electropores created at the elevated temperature. Indeed, the viscosity of *E. coli* phospholipids changes with temperature (Renne and Ernst 2023; Sinensky 1974), and for other cell types, it has been shown that membrane elasticity and fluidity is significantly increased when the temperature is increased from 4 °C to 30 °C, leading to considerably lower membrane breakdown voltage at higher temperatures (Coster and Zimmermann 1975; Kandušer, Šentjurc, and Miklavčič 2008). From these observations, it follows that electropores can more easily be created at 28 °C, compared to at 6 °C, and optimization of pulse parameters should be explored further to search for a potential compromise between limited membrane damage and sufficient delivery of desired molecules at elevated temperature.

### 2.3. Off-chip electroporation with bipolar pulse

Given the detectable difference in electroporation results obtained with monopolar and bipolar square pulses in the microfluidic system, we further investigated if this difference was also observable in bulk conditions. Normally, a short, high-voltage pulse for bulk electroporation is created by discharging a capacitor, which produces exponentially decaying voltage. To the best of our knowledge, square bipolar pulses have not been reported before for bacteria electroporation, most likely due to the complexity of the electrical circuit required to obtain a kV-amplitude ms-long pulse of alternating polarity. To produce such a pulse, we used a setup similar to the on-chip electroporation device but with a different high-frequency amplifier capable of delivering up to a few hundred volts.

We aimed to perform bulk electroporation of the same cell strain under conditions as similar as possible to the on-chip electroporation experiments. In brief, cells were grown in bulk liquid culture at 37 ⁰C, cooled on ice, and washed with 10 % glycerol by exchanging the medium in multiple rounds of cell sedimentation by centrifugation. We found that using pure deionized water without glycerol for cell washing led to high cell mortality even without pulsing. We speculate that this is due to a combination of increased osmotic pressure in pure water and the mechanical force applied during centrifugation. Thus, 10 % glycerol was used in all bulk electroporation experiments reported below. Cy5-labeled DNA oligonucleotides were added to the cell suspension which was then transferred to a custom-made aluminum electroporation cuvette with a 0.56 mm gap between the electrodes. The cell-containing cuvette was pulsed with a 1 ms square monopolar or bipolar pulse of 616 V, which resulted in an electric field of 1.1 V/μm, similar to the electric field in the microfluidic device. After the pulse, cells were rapidly diluted in growth medium, sparsely placed on an agarose pad, and finally imaged under the microscope to monitor electroporation efficiency and cell growth (Figure S6). The single-cell data was analyzed in the same way as the on-chip data and cells were classified into the same four classes as before (Figure 2f).

We found that monopolar and bipolar 1 ms pulses resulted in very similar results: ∼51-57 % of “green” cells, ∼3-4 % of “yellow” cells, ∼5 % of “red” cells, and ∼35-40 % of “grey” cells (Figure 9). The difference in outcome between monopolar and bipolar pulses in bulk is within experimental error, as opposed to the on-chip results, where the bipolar pulse resulted in a somewhat higher fraction of Cy5-DNA permeabilized cells and consequently higher fraction of “green” cells. The fraction of dead cells (“red” and “yellow”) was also much lower after being pulsed in bulk compared to on-chip, and the fraction of detected electroporated live cells (“green”) was much higher (∼60 % maximum in bulk *vs*. ∼15 % maximum on-chip). We note that differences in apparent permeabilization efficiencies might be partially explained by the improved signal-to-background ratio for Cy5-DNA detection on an agarose pad compared to the microfluidic device. That is, some Cy5 dye might be irreversibly and non-specifically adsorbed to the PDMS surface even after liquid exchange and washing, resulting in a higher fluorescence background. Also, we cannot exclude that some highly damaged cells were completely disintegrated before deposition onto the agarose pad, leading to “survivorship” bias, *i.e.* underestimation of dead cells for the bulk experiment.

**Figure 9.**
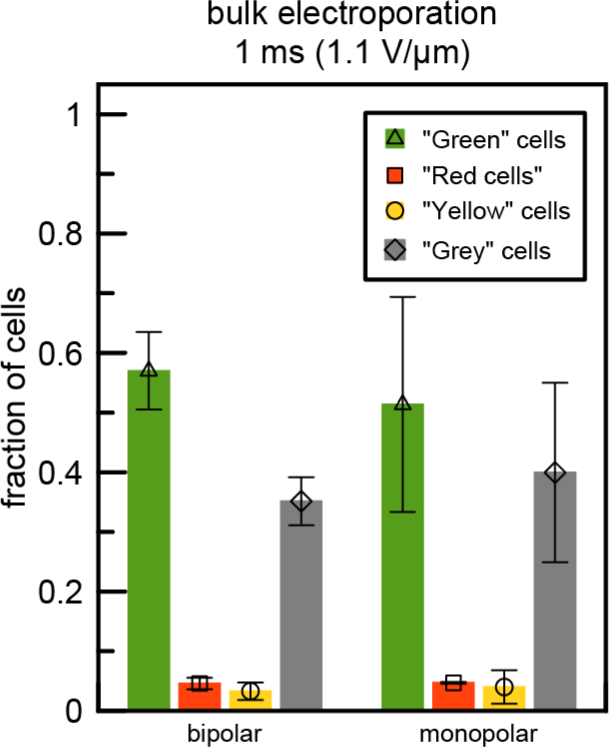
Results of bulk electroporation with square monopolar and bipolar pulses at 1.1 V/μm field strength. Cell classifications are described in Figure 2f. Error bars represent standard deviation derived from two independent experiments.

The exact reasons for the observed differences in cell viability and internalization efficiency in bulk *vs.* on-chip experiments with the same type of electrical pulse are not clear. We speculate that they could originate from differences in environmental parameters (10 % glycerol *vs.* deionized water, 0 ⁰C *vs.* 6 ⁰C), geometrical parameters (cells in random orientation *vs.* pre-aligned to the electrical field, absence *vs.* presence of cell-to-cell contacts), and/or mechanical parameters (free electrophoretic movement *vs.* mechanically restricted movement and collisions) although a focused specific study would be required to identify the precise mechanisms. As a bipolar pulse does not improve permeabilization nor cell viability, we hypothesize that under our experimental conditions, the pulse does not change the properties of the formed electropores in the cell membrane. Instead, its positive effect observed in the microfluidic device stems from limiting the negative effects related to DNA and cell electrophoresis which are pronounced on-chip when the distance between the electrodes is on the same length scale as the electrophoretic distance.

## 3. Conclusion

In this work, we designed, fabricated, and used a microfluidic platform for a systematic *in situ* study of electroporation of bacteria. As a proof-of-concept, the delivery of short, fluorescently labeled nucleic acids in *E. coli* cells was demonstrated, showing the potential for electroporation of other bacteria and delivery of different types of molecules, such as plasmid DNA and proteins. With some design modifications, this platform can be transformed into a high-throughput single-cell electroporation device. Based on the present study, we can highlight some important considerations when designing such a microfluidic device for reversible electroporation of bacteria.

We discovered that electrophoresis of cells and macromolecules plays a critical role when performing electroporation in small volumes. The effect causes a decrease in the local concentration of DNA between electrodes and movement of cells during the pulse away from the optimal position between electrodes. To counteract these effects, we propose to use a bipolar electroporation pulse, which in our setup optimized the yield of live cells with internalized molecules (“green” cells). Among the tested parameters, a 1 ms, 1.1 V/µm bipolar pulse resulted in the highest fraction of live and electroporated cells (∼15 %). This efficiency is significantly lower than that of cuvette electroporation under similar pulse conditions (∼60 %), but, considering the advantage of a controlled cell microenvironment and compatibility with *in situ* microscopy, the efficiency is still, in our opinion, high enough for the system to be exploited for single-cell biological studies including single-particle tracking of delivered fluorescent molecules.

Further, we found that when the space between cells and electrodes is filled with highly resistant liquid, a voltage divider effect occurs. This considerably reduces the field strength across the cells and thus decreases the probability of electroporation. Therefore, the position of the cells with respect to the electrodes should be precise with a minimal gap between the electrodes and the cells.

Another important observation from our study is that cell cooling before and during the pulse is essential for cell viability. We suggest that this originates from a temperature-dependent change in the physical properties of the cell membrane, rather than harmful Joule heating produced by the electrical current. However, we note that this only holds if the resistance of the surrounding liquid is high (electrical current is low) and that it cannot be directly extrapolated to bacterial species other than *E. coli* due to possible differences in cell physiology.

Finally, we conclude that regardless of precisely controlled environmental conditions and geometrical parameters of the microchip, electroporation results in a highly heterogeneous outcome with respect to cell survival and internalization of DNA into the cells. Although we cannot completely rule out that this heterogeneity is caused by imperfections in the manufacturing process of our miniaturized platform, it likely originates from naturally occurring cell-to-cell variations in the cell membrane composition (Ginez et al. 2022), cell cycle stage at the time of the pulse, variation in gene expression, and the stochastic nature of the stability and number of electropores formed (Glaser et al. 1988; Kotnik et al. 2019). This observation further underscores the importance of a platform capable of performing single-cell electroporation, both for the direct study of the process to better understand the factors responsible for the cell-to-cell variations in electroporation outcome, and for advanced single-cell studies of electroporated cells.

## 4. Experimental Section

### Fabrication of the microfluidic electroporation device

The microfluidic device was composed of polydimethylsiloxane (PDMS, Sylgard 184) microfluidic channels bonded to a 175 µm thin glass (Borofloat 33) forming the bottom of the microfluidic channel with patterned electrodes. The PDMS channels were prepared by molding the PDMS pre-polymer solution on a silicon master patterned by a combined electron-beam and UV lithography process followed by dry etching of silicon dioxide, as previously described (Baltekin et al. 2017). The channels have a cross-section of 1.2 *x* 1.2 µm^2^ and a length of 50 µm with a 300 nm constriction at the end to ensure cell trapping in the channels. During PDMS molding, stainless steel tubes with an outer diameter of 0.8 mm were placed 400 µm above the silicon master to form cooling channels in the PDMS chip body, following the procedure described elsewhere (Khaji and Tenje 2022). The electrodes were patterned on the glass bottom of the microfluidic channels by sputtering 20 nm of tantalum as the adhesion layer, followed by 110 nm of platinum onto a UV lithography patterned resist mask (1813, Shipley), which was afterwards removed by a lift-off step in Acetone. Once the electrodes were patterned, the glass wafer was diced into 8 chips and individually bonded to the PDMS microfluidic channels by placing them in contact for at least 1 h at 100 °C by carefully aligning the two pieces after an air plasma treatment of the PDMS surface for 30 s (D-50E Heavy Duty Generator).

### Microfluidic setup

Experiments were performed under continuous perfusion of liquids through the microfluidic chip. The PDMS chip layout is shown in Figure S1. Liquid reservoirs (15 ml polypropylene tube containing 3‒6 ml of liquid) were pressurized using a pressure controlling unit (OB1 MK3+, Elveflow) connected to the microfluidic chip by Tygon tubing (0.51 mm ID) according to the scheme shown in Figure 1. Tubings from cell media and water/Cy5-DNA reservoirs were connected to the chip via remotely controlled 2-way liquid valves (Elveflow). Washing of the back channels was driven by gravity from a reservoir with the liquid level 7 cm above the chip level.

The cooling system of the microchip comprised two polyetheretherketone (PEEK) tubings (0.5 mm ID, 0.79 mm OD) inserted in the PDMS chip 400 µm above the microfluidic channels (Khaji and Tenje 2022), connected with silicone and polytetrafluoroethylene (PTFE) tubings (1 mm ID, ∼70 cm length) to two 2 l liquid reservoirs pressurized by a pressure controlling unit (OB1 MK3+, Elveflow). The reservoirs were placed on ice and filled with a water-ice mixture before the experiment to supply 0°C water to the chip. When activated, the flow rate through the cooling channel was 18 ml/min which resulted in chip temperature of 6°C.

### Electrical circuit of the microfluidic device

Four electrode blocks (two with 20 μm distance between the electrodes, two with 16, 18, 20 and 22 μm stepwise variable distance between electrodes) were overlaid with the microfluidic cell traps (Figure 1 and S1). The chip was glued with silver-epoxy (CircuitWorks, Chemtronics) to a PCB interface comprising contact wires for the external equipment necessary to perform the electroporation. A function generator (AFG3022C, Tektronix), connected to a high frequency amplifier (WMA-300, Falco Systems) was used to supply the electrical pulse. As default, all electrode blocks were grounded via a ground bridge wire on the bread-board. Upon electroporation, the ground bridge was manually removed from that specific block of electrodes a few seconds prior to the pulse and bridged back a few seconds after the pulse. Amplified voltage was applied to the desired set of electrodes connected in series with a 10 kΩ resistor. One channel of an oscilloscope (GDS-1052-U, GW Instek) was connected to the amplifier output terminal via a 10X probe to monitor the voltage, and the second channel was connected to the10 kΩ resistor via a 1X probe to measure voltage drop on the resistor and detect the current passing through the circuit. The resistance of the chip (measured independently) was ∼1MΩ, and thus, ∼99 % of the voltage reached the electrodes of the chip.

### Electrical setup for bulk electroporation

Electroporation cuvettes were made in-house from two rectangular aluminum blocks (SK 480 25, Fischer Elektronik) covered with aluminum conductive sticky tape (1436, 3M) to which two copper wires were soldered. The 0.56 mm gap between the blocks was made using two layers of non-conductive sticky frames (AB0578, Thermo Fisher Scientific). The electrical pulse was generated using a function generator (AFG3022C, Tektronix) connected to a high frequency piezo driver/amplifier (PZD700A, Trek). The voltage applied to the cuvette and current through the circuit were measured directly from monitor outputs of the amplifier by a two-channel oscilloscope (GDS-1052-U, GW Instek).

### Optical setups

Imaging of the microfluidic electroporation process was performed on a laser scanning confocal microscope (SP8, Leica) equipped with a 20x air objective (HC PL FLUOTAR 20x/0.50), photomultipliers and a hybrid technology detector (HyD), as well as 488 nm and 638 nm lasers. Bright field images were acquired by detecting transmitted light under 488 nm illumination while the fluorescence Cy5 signal was detected under 638 nm excitation.

Imaging of agarose pads was performed on a widefield epifluorescence microscope (Nikon Eclipse Ti2) with a CFI Plan Apo DM lambda 1.45/100x oil objective (Nikon). Phase contrast and fluorescence images were recorded with an ORCA-Quest qCMOS camera (Hamamatsu Photonics) connected through an additional demagnifying 0.7x camera adapter (Nikon). Fluorescence images were acquired with 642 nm laser excitation (MPB Communications) at 10 ms continuous exposure of 130 W/cm^2^ power density using an in-house-written µManager (Edelstein et al. 2010) plug-in.

### Growth media, liquids, and cell strain

Experiments were performed with the *E. coli* MG1655 strain. Super Optimal Broth (SOB) media (2 % (w/v) tryptone, 0.5 % (w/v) yeast extract, 10 mM NaCl, 2.5 mM KCl, 20 mM MgSO_4_) was used for over-night cultures. Where applicable, the SOB media was supplemented with 85 μg/ml of surfactant (Pluronic F-127, Sigma-Aldrich) and 1 μM SYTOX Blue dead-cell stain (Invitrogen). The surfactant was necessary to prevent bacteria sticking on the PDMS surface, and SYTOX Blue was used for quick assessment of cell viability during the experiments (this assessment was not used in data analysis). Deionized water (resistivity 18.2 MΩ·cm at 25 °C, Milli-Q) containing 0.4 µM 3’-Cy5-labeled DNA oligo (GGGAGATCAGGATA, HPLC-purified, Integrated DNA Technologies) was used for cell washing before and during electroporation. All liquids were filtered with 0.2 µm syringe filters before use.

### Experiment workflow for on-chip electroporation

Cells from over-night culture (grown at 37° C in SOB) were diluted 2,000 times in SOB with surfactant. After a few hours of growth at 24±2 °C, exponentially growing cells (OD_600_ < 0.1) were loaded into the microfluidic chip. Cell loading was typically achieved in 20-30 min, after which the cooling system was activated and left to stabilize for 5 min. “Pre-pulse” images of the cells on the chip were acquired at this stage, and 25 min after cooling had started the growth media perfused through the chip (SOB with surfactant and SYTOX Blue) was changed to deionized water containing Cy5-DNA by switching the remotely-controlled valves (Supplementary Video 1). Delivery of liquid to the front channel of the chip, the cell traps, as well as back channels and connected liquid channels, was controlled by following the Cy5 fluorescence signal in the microscope. After 10 min of water/Cy5-DNA wash, the grounding bridge from the pulsing electrodes was removed, and an electroporating pulse was applied to one of the four electroporation blocks, while simultaneously recording fluorescence and bright field movies of the pulsed region (at 19 ms/frame, or 56 ms/frame for pulses of reversed polarity). The grounding bridge was returned, and 40 s after the pulse SOB with surfactant and SYTOX Blue was supplied to the cells instead of water/Cy5-DNA. The cooling system was stopped 3 min after the pulse when all water in the cell traps had been replaced with growth media. “Post-pulse” images of cells were acquired 10 min after switching to growth media, and cell growth was monitored in total 60 min after the pulse with 5 min intervals between image frames.

### Experiment workflow for bulk electroporation

Cells from over-night culture (grown at 37 °C in SOB) were diluted 200 times in SOB with surfactant and SYTOX Blue. After a few hours of growth at 37 °C, exponentially growing cells (OD_600_ = 0.2) were placed in an ice-water bath for 20 min. All subsequent steps were performed in the cold room (4 °C). Cells were pelleted by centrifugation in 50 ml Falcon tubes (2,000 × g, 10 min). The supernatant was removed and cells were resuspended in the same 50 ml Falcon tubes in 10 % (v/v) glycerol/water solution using cut pipette tips (providing bigger holes and thus less disturbance for cells) and pelleted again by centrifugation (1,500 × g, 10 min). The pelleted cells were then washed additionally four times by centrifugation (1,000 × g, 5 min) in 2 ml tubes. After the final wash, the cells were resuspended in 10 % (v/v) glycerol and the cell density was adjusted to OD_600_ = 100. A test electroporation of 5 µl of pure cell suspension was performed (1 ms monopolar square pulse, 1.1 kV/mm), which typically resulted in 21 ± 2 kΩ resistance in the sample (calculated from the current monitor signal of the amplifier, equivalent to 33 ± 3 kΩ·cm or 3 ± 0.3 mS/m). A similar test pulse of pure 10 % (v/v) glycerol solution resulted in similar resistance (22 kΩ), showing that the established washing procedure efficiently removed salts from the cell media. Cells were mixed with Cy5-labeled DNA oligo (0.4 µM final concentration). 5 µl of the cell suspension was placed between the electrodes of an ice-cold electroporation cuvette and pulsed 65 s after the cuvette was removed from ice to the lab bench. Cells were immediately removed from the cuvette, diluted in SOB to OD_600_ = 0.1 and sparsely placed on a 2 % agarose pad. The agarose pad was prepared with SeaPlaque GTG Agarose (Lonza) and SOB supplemented with surfactant and SYTOX Blue. The pad was sealed between the microscope slide and the coverslip (#1.5H, Thorlabs) using a sticky plastic frame (AB0578, Thermo Fisher Scientific), and then mounted on the microscope, where single cells were imaged at 28 ± 2°C. Acquisition of phase contrast images and Cy5 fluorescence was made every 30 min for 1 h starting ∼1 h after electroporation.

### Data analysis

Cy5 fluorescence, bright field (for microchip imaging), or phase contrast (for agarose pad imaging) images were processed with custom scripts in MATLAB and then annotated manually. In brief, the Cy5 signal was converted to a binary mask (Figure 2b (inset)) and overlaid with cell images (Figure 2d). Images of cells acquired sequentially were combined in one multi-frame file with automatic drift correction of individual frames. Multi-frame files containing the Cy5-mask and images of growing cells were analyzed using the “Cell Counter” ImageJ plugin by manual annotation of each cell into one of the four classes depending on visible cell growth and visible cell mask overlapping with the cell image on the first frame (Figure 2e, f). For on-chip experiments, cell traps were classified as “overfilled”, “normal” and “underfilled” according to Figure 3a and only cells from “normal” traps were taken for the analysis, unless otherwise stated. When a cell on the agarose pad had an excessively high Cy5 signal, possibly masking the Cy5 signal of the neighboring cells (bulk experiments), or a cell was not visible behind the electrodes (on-chip experiments), classification was not possible and, thus, these cells were not included in the analysis (∼20 %). Typically, for each set of experimental conditions (*e.g.* field strength or pulse duration), the datasets from on-chip and bulk electroporation included 50-200 and 2,000-6,000 cells, respectively. For the bulk electroporation, the error estimates (error bars in Figure 9) were computed as a standard deviation from two independent experiments. Reproducibility errors of the on-chip experiments were derived by computing the coefficient of variation (ratio of the standard deviation to the mean) for data points obtained under close field strengths (1.091, 1.1 and 1.125 V/µm) at 2 ms bipolar pulses in three independent experiments (Figure S5). The data were approximated by step function *CoV(x)* = [0.5 if *x* < 0.2; 0.3 if *x* ≥ 0.2]. Thus, the reproducibility error for each data point (*p_i_*, fraction of cells of certain class *i*) was computed as *E_rep i_* = *Cov* · *p_i_*. To estimate the measurement error caused by limited sample size for individual on-chip experiments, we assumed that cell classification counts follow a multinomial distribution, for which the expected number of cells of class *i* counted for any given experimental condition is *x_i_* = *np_i_*, where the total number of cells counted for each given experimental condition is *n*, and the fraction of cells of a certain class *i* is *p_i_*. Then, the variance of *x_i_* is *Var*(*x_i_* = *np_i_*(1 − *p_i_*), and the standard deviation of *p_i_* is given by the formula: 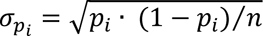. The obtained errors *E_rep i_* and *σ_pi_* were combined as 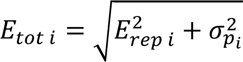 and plotted as the error bars in Figures 3, 4, 6, 7 and 8.

## Data and software code availability

Experimental data generated and analyzed during the current study and image analysis code are available in SciLifeLab Data Repository (https://doi.org/10.17044/scilifelab.25966867).

## Acknowledgements

This work was funded by the Swedish Research Council (M.J. and M.T.: 2016-06213, M.J.: 2019-03714), the European Research Council (M.J.: SMACK-947747) and SciLifeLab via a postdoc grant to ZK. We acknowledge Myfab Uppsala for providing facilities and experimental support. Myfab is funded by the Swedish Research Council (2019-00207) as a national research infrastructure. We are grateful for help from Dr. Praneeth Karempudi, Uppsala University with preparation of the Si mold used for PDMS chips and Dr. Irmeli Barkefors and Dr. Susan Peacock, both Uppsala University, for careful proof-reading of the manuscript.

