## Supplementary Information for "A microfluidic platform for *in situ* studies of bacteria electroporation"

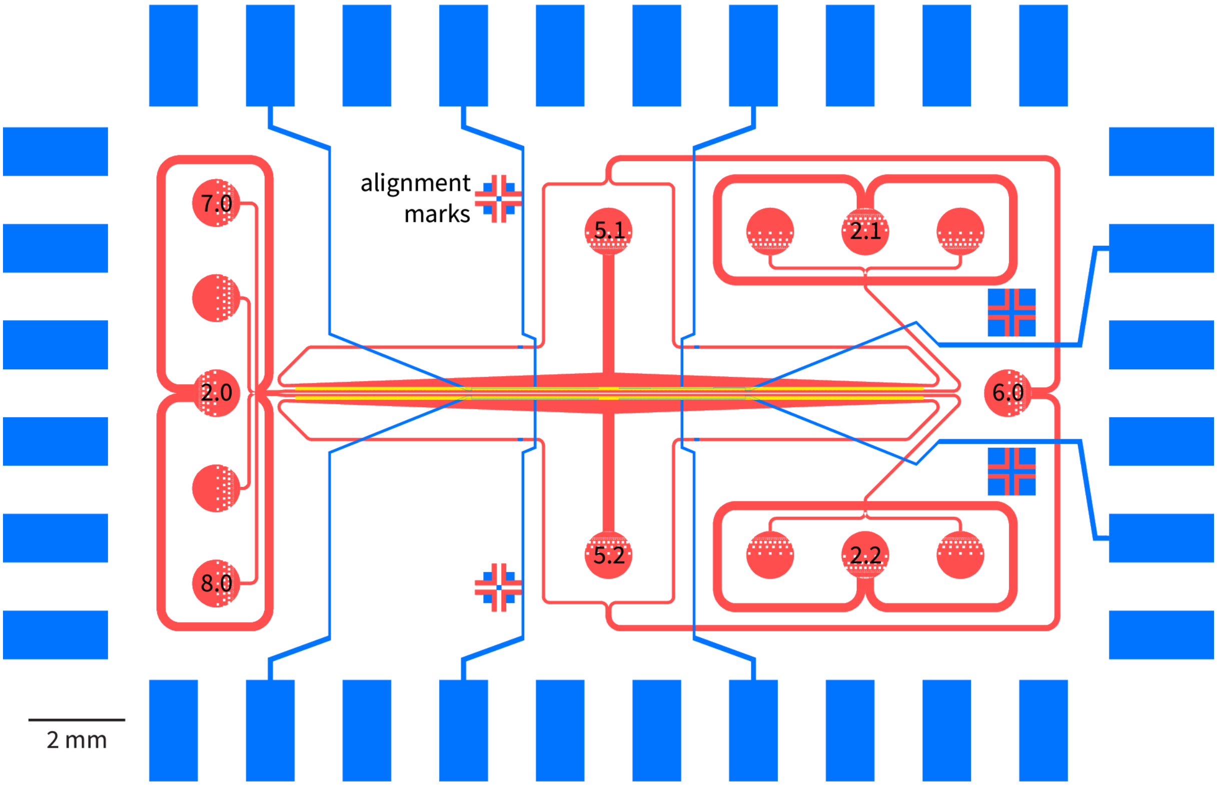


**Figure S1.** Drawing of the chip with microfluidic channels (red), cell trap regions (yellow) and platinum electrodes (blue). Numbered liquid ports were punched and used as follow: 2.0 – waste outlet, 2.1 and 2.2 – cell loading (waste when loading was done), 5.1 and 5.2 – pure deionized water, 6.0 – waste, 7.0 – cell media, 8.0 – water/Cy5-DNA.

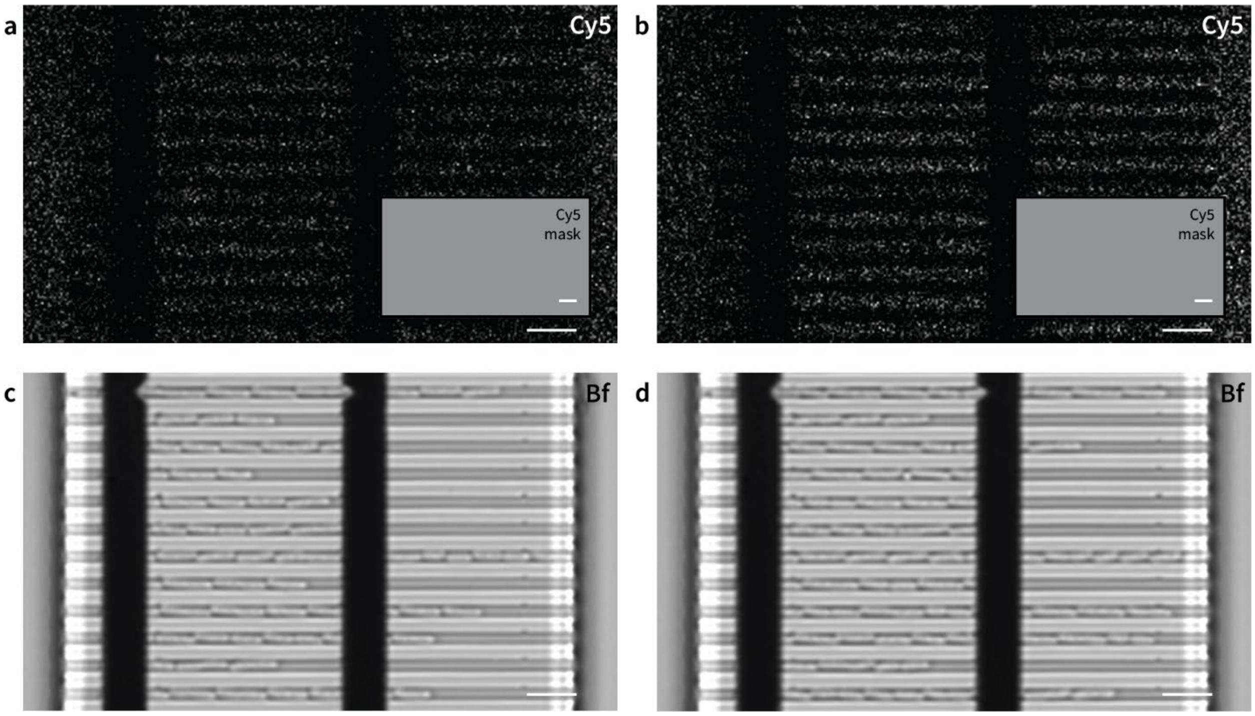


**Figure S2.** Negative control for electroporation procedure. Cells loaded between grounded electrodes and imaged before (a, c) and after (b, d) electroporation performed on a neighboring electrode block on the same chip. Cy5-fluorescence (a, b), Cy5-fluorescence mask (inset in (a, b)) and bright field images, overlaid with white Cy5-fluorescence mask (c, d). Scale bars (in white) are 5 µm. Corresponding cell growth time-lapse shown in Supplementary Video 2.

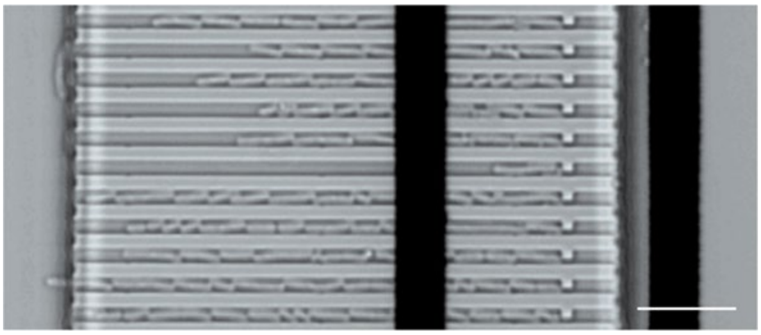

**Figure S3.** Example bright field image of the electrodes block misaligned relative to cell traps. Scale bar (in white) is 10 µm.

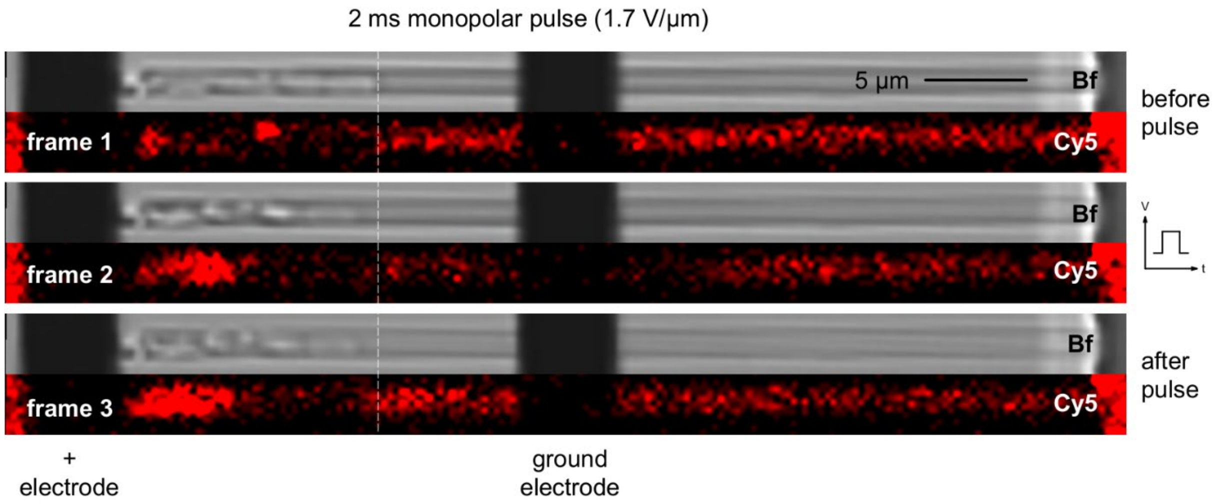


**Figure S4.** *In situ* observation of electroporation with 2 ms 1.7 V/µm square monopolar pulse of reversed polarity (relative to the experiment shown in Figure 5a) recorded at 56 ms per frame with bright field and Cy5-fluorescence imaging. Ground and pulsed electrodes are marked. Cells move from the ground to the positive electrode. White dashed line shows the position of the cell pole before the pulse.

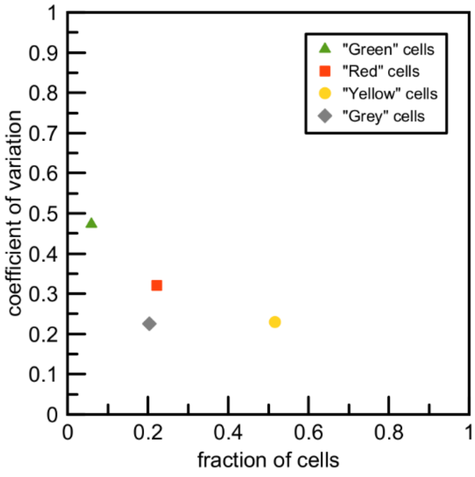


**Figure S5.** Coefficient of variation (the standard deviation over the mean) computed for data from 3 independent electroporation experiments performed under similar conditions (2 ms bipolar pulse at 1.091, 1.1 and 1.125 V/µm, datapoints from Figure 4f).

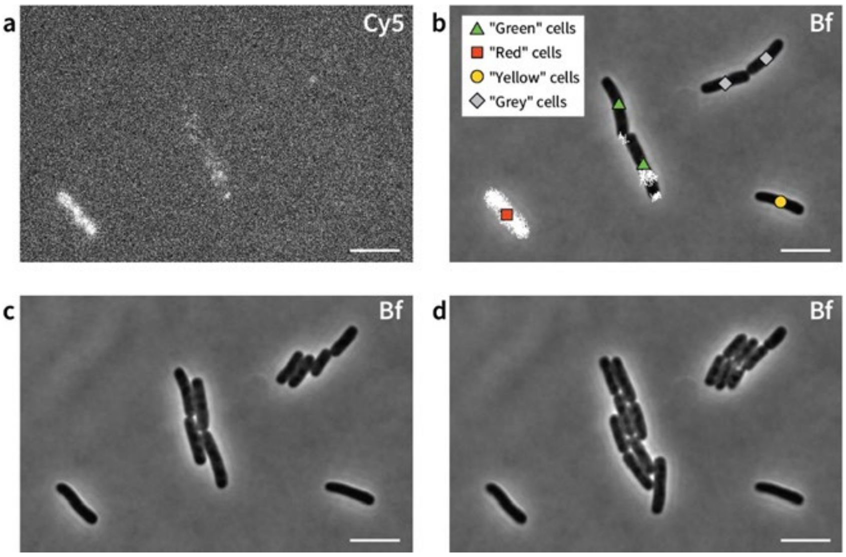


**Figure S6.** Microphotographs of cells electroporated in cuvette with 1 ms 1.1 V/µm square bipolar pulse and imaged on an agarose pad. Cy5-fluoresence (a) and phase contrast (b-d) images acquired immediately after pad preparation (a, b), 30 min later (c) and 60 min later (d). On (b) phase contrast image is overlaid with white Cy5-fluorescence mask and cells are classified according to Figure 2f. Scale bars (in white) are 5 µm.

### Supplementary Note 1

**Conductivity of water containing Cy5-DNA.** A concentration of 0.4 µM of 14-mer Cy5-DNA is equivalent to 14·0.4 µM = 19.6 µM of DNA monomers. Though we cannot find reference values for molar conductivity of DNA, we assume that it does not exceed the conductivity of inorganic salts. These have molar conductivity $\Lambda_m$ in the order of ~10 $S\cdot M^{-1}m^{-1}$ (Lide 1992). Hence, at 19.6 µM concentration, the conductivity of the DNA solution should not exceed $2\cdot10^{-4}$ $S\cdot m^{-1}$ = 0.2 $mS\cdot m^{-1}$.

In our experiments we used deionized water (Milli-Q) with resistivity ρ = 18.2 MΩ·cm at 25°C. This is equivalent to conductivity $\sigma = 1/\rho = 5.5\ 10^{-8}\ S\cdot cm^{-1} = 5.5\ 10^{-6}\ S\cdot m^{-1}$, which is negligible compared to the conductivity contributed by DNA.

### Supplementary Note 2

**Calculation of the temperature rise due to Joule heating effect on the chip.** In our calculations we assume that Joule heating produced by the current between electrodes during the pulse is dissipated in the volume of the microchannels between electrodes. The temperature rise is calculated from known parameters of the typical pulse, chip geometry, and properties of the heating material. We note that this is an upper bound value since it does not account for energy dissipation during the pulse, and heat transfer due to constant liquid exchange on the chip. We made two separate calculations assuming that the whole volume is filled with only water or that the whole volume is occupied by cells (Table S1, see Table_S1.xlsx Supplementary File). In both cases, the estimated temperature increase is ~4-5°C.

**Supplementary video legends**

**Supplementary Video 1.** Region of the chip between the blocks of electrodes during exchange of cell media to water/Cy5-DNA solution imaged in bright field (grey) and Cy5-fluorescence (red) every 1 s. Playback speed is 1 frame per second (*i.e.* real time).

**Supplementary Video 2.** Cell growth in negative control for electroporation procedure. Cells loaded between grounded electrodes imaged in bright field every 5 min after electroporation procedure performed on the same chip but a neighboring electrode block. Playback speed is 3 frames per second (*i.e.* 900 times faster than reality). Corresponding Cy5 and bright field images acquired before and after electroporation procedure are shown in Figure S2.

**Supplementary Video 3.** Cell growth imaged every 5 min in bright field after electroporation with a 1 ms 1.2 V/µm monopolar pulse. Playback speed is 3 frames per second (*i.e.* 900 times faster than reality). Corresponding Cy5 and bright field images acquired before and after electroporation procedure are shown in Figure 2a-d.

**Supplementary Video 4.** *In situ* observations of electroporation with 2 ms square monopolar electrical pulse of 1.4 kV/µm with images recorded every 19 ms in bright field (upper) and Cy5-fluorescence (lower). Playback speed is 3 frames per second (*i.e.* 17.5 times slower than reality).

**Supplementary Video 5.** *In situ* observation of electroporation of 2 ms square bipolar electrical pulse of 1.33 kV/µm with images recorded every 19 ms in bright field (upper) and Cy5-fluorescence (lower) imaging. Playback speed is 3 frames per second (*i.e.* 17.5 times slower than reality).
